# Quantitative prediction of enhancer-promoter interactions

**DOI:** 10.1101/541011

**Authors:** P.S. Belokopytova, E.A. Mozheiko, M.A. Nuriddinov, D. Fishman, V. Fishman

**Author notes:** correspondence should be addressed to Veniamin Fishman.

## Abstract

Recent experimental and computational efforts provided large datasets describing 3-dimensional organization of mouse and human genomes and showed interconnection between expression profile, epigenetic status and spatial interactions of loci. These interconnections were utilized to infer spatial organization of chromatin, including enhancer-promoter contacts, from 1-dimensional epigenetic marks. Here we showed that predictive power of some of these algorithms is overestimated due to peculiar properties of biological data. We proposed an alternative approach, which gives high-quality predictions of chromatin interactions using only information about gene expression and CTCF-binding. Using multiple metrics, we confirmed that our algorithm could efficiently predict 3-dimensional architecture of normal and rearranged genomes.

## Introduction

Spatial interactions between promoters and their regulatory sequences are required to maintain cell-type specific expression pattern ^1^. Intriguingly, enhancers do not necessarily regulate closest promoters, and enhancer-promoter (EP) interactions often span large genomic distances ^1,2^. Although enhancer targets can be directly identified using high-resolution 3C-methods ^1^, these data is costly and currently available only for a small subset of cell types. Besides, experimental identification of enhancer targets does not provide mechanism explaining target selection.

To address these challenges, several computational tools were developed, aiming to predict 3-dimensional EP interactions from 1-dimensional genetic and epigenetic marks ^3–12^. All these tools utilize one of the two approaches. First type of tools relay on building a physical model of chromatin, and optimize model parameters so that it fits best experimental (usually Hi-C) data ^7,10^. The optimized model can be used to infer spatial conformation of chromatin, including those regions containing EPI. Another approach is to find consistent patterns in epigenetic data which would explain 3-dimensional contacts of chromatin ^3–6,8,9,11,12^ This approach is promising, since it can, in theory, capture dependencies between epigenetic states of the chromatin and its genomic organization even without complete *a priori* knowledge of physical bases underlying 3-dimensional organization of chromatin.

Here we aimed to develop an algorithm for prediction of Enhancer-promoter interactions. We first benchmarked existing *TargetFinder* algorithm ^5^, which is of particular interest, because of its high accuracy, low false-discovery rate and reproducibility, demonstrated by analysis of several human cell types. Surprisingly, we found that efficiency of this and similar algorithms was strongly overestimated due to incorrect design of training and validation datasets. Moreover, we showed that EP interactions are quantitative rather than qualitative, and designed a novel approach to estimate EP interactions based on epigenetic properties of loci. We showed that information about gene expression and CTCF-binding is sufficient for accurate prediction of EP interactions, and validated this model on several experimental human and mouse datasets. Finally, we demonstrated that our model can be utilized to predict alterations of 3-dimensional genome organization caused by chromosomal rearrangements.

## Results

### *TargetFinder* fails to predict EP interactions

We have started our research from extending existing *TargetFinder* algorithm for prediction of EP interactions in mouse data. Originally, *TargetFinder* was developed and validated on human datasets. We annotated promoters and enhancers as interacting and non-interacting using hi-resolution Hi-C data on mouse ES cells ^13^ and collected a set of 24 genetic and epigenetic predictors (Table 1). To construct our datasets, we used original definition of “interacting” promoters and enhances proposed in *TargetFinder* paper ^5^, i.e. promoter and enhancer were considered as interacting only if they were located in the anchors of Hi-C loop. Surprisingly, *TargetFinder* accuracy (measured by either precision, recall or f1-score) was dramatically lower than previously reported on human data (Table 1). We found that changing ratio of interacting to non-interacting EP pairs from 1:20 to 1:1 increases f1-scores; however, obtained values were still far below than reported previously ^5^ (Table 1). We additionally run *TargetFinder* on mouse cortex and neural stem cells (NPC) ^13^ using 10 available epigenetic predictors. Similarly to results obtained on ES cells data, *TargetFinder* was not efficient on these datasets (Table 1).

**Table 1.**

| Cell type | Number of predictors | Train and validation datasets | F1-score |  |
| --- | --- | --- | --- | --- |
|  |  |  | Interacting : Non-interacting<br>1:20 | Interacting : Non-interacting<br>1:1 |
| Mouse ES cells | 24 | This paper | 0.023391812 | 0.674285714 |
| Mouse cortex | 10 | This paper | 0.081967213 | 0.7529411764 |
| Mouse NPC | 10 | This paper | 0.123893805 | 0.735202312 |
| Human GM12878 | 100 | This paper | 0.07 | 0.5897435897 |
| Human GM12878 | 100 | Original | 0.83 |  |

To understand why *TargetFinder* fails to predict EP interactions we re-processed original human data generating predictors, training and validation datasets for human GM12878 cells *de novo*. Surprisingly, running *TargetFinder* on these re-processed human datasets resulted in low f1-scores, with only small improvement comparing to mouse ES cells data (Table 1).

Comparing our protocol of data processing with the pipeline that was used to generate original *TargetFinder* datasets we noticed the difference in composition of training and validation samples. In the original approach EP pairs were randomly split to obtain training (∼90% of data) and validating (∼10% of data) datasets. Our pipeline randomly selects 2 chromosomes and designs all EP pairs on these chromosomes as validation dataset (∼10% of all data), and the rest of EP pairs as training dataset. The difference between approaches appears when genomic region between promoter and enhancer of a EP pair (referred hereinafter as “window”) overlaps a window of another EP pair (Fig. 1, A). This is the case, for example, when single promoter interacts with several enhancers, thus forming multiple EP pairs with shared promoter- and window regions (of note, 69% of all promoters in GM12878 dataset interact with several enhancers). In the original *TagetFinder* pipeline, EP pairs with the overlapping windows were randomly distributed between training and validation datasets. In contrast, our re-processed dataset would never contain pairs with windows overlapping between training and validation samples.

**Figure 1.**
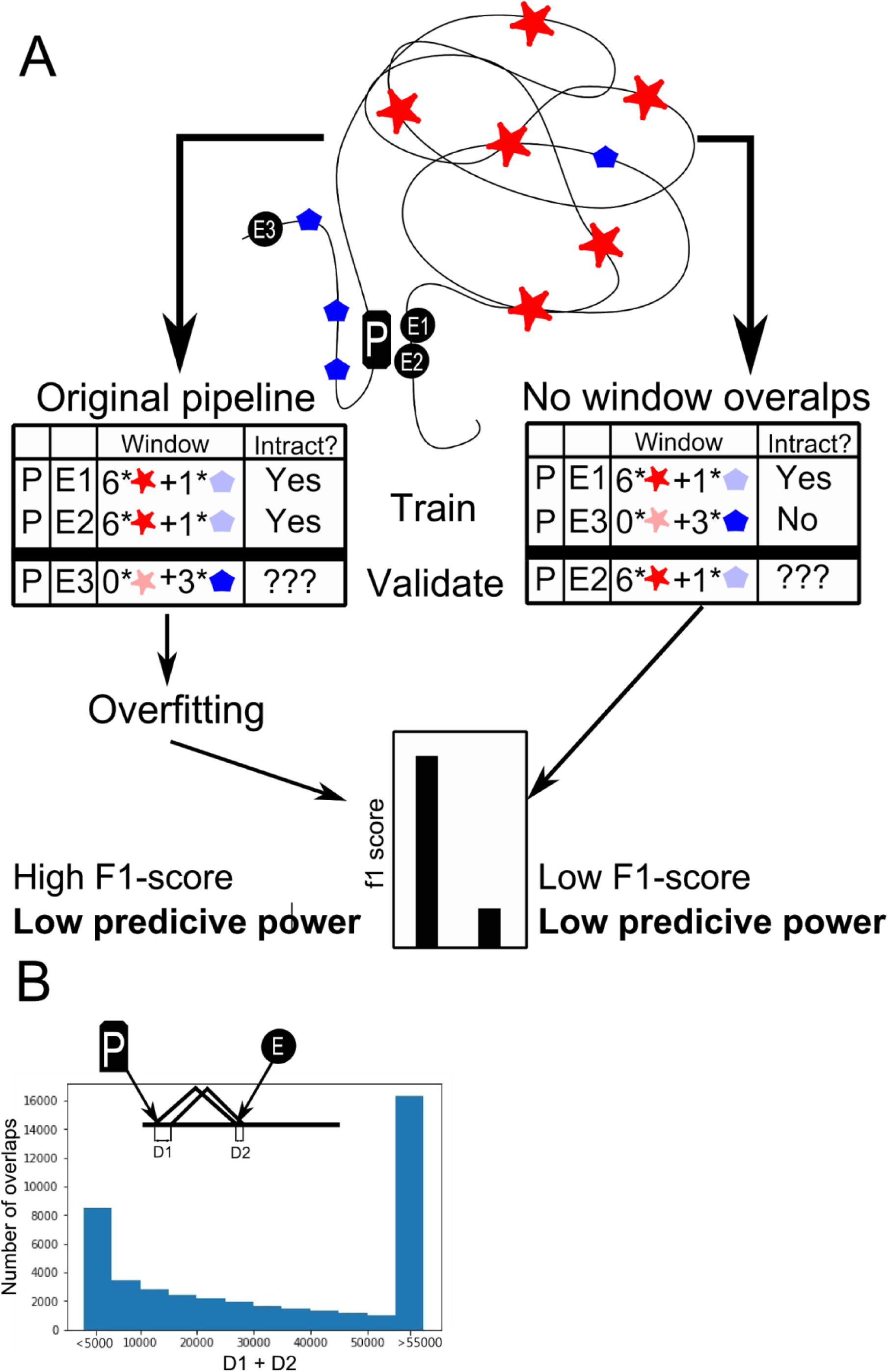
**A.** Schematic illustration of the original and corrected *TargetFinder* pipelines. **B**. Distribution of distances between borders of overlapping EP pair windows. For each EP pair we found the window of another EP pair so that distance between borders of windows (*d=D1+D2*) is minimal. Histogram shows distribution of the obtained *d* values.

One should note, that overlaps of EP windows are frequent (>99% of all pairs share window with at least one another pair) and overlap size is often large (Fig. 1, B), which means that promoters and enhancers sampled from such pairs are located close to each other in genome. Since spatial contacts of neighboring genomic regions are correlative, interactions of a EP pair should be a good predictor for interaction of another EP pair when both promoters and enhancers are located nearby (i.e. if window overlap is high). To confirm this, for each EP pair we find another pair from GM12878 dataset with highest window overlap and used interaction of the latter pair as a predictor of interaction of the former pair. The prediction was almost perfect, characterized by f1-score ∼0.9. Finally, we note that large overlap of windows of two EP pairs would result in similar, if not identical, values of predictors within these windows, which allows identification of highly overlapping pairs in absence of information about genomic coordinates of promoters and enhancers.

Considering that *TargetFinder* efficiency drops dramatically when regions shared between training and validating datasets are excluded, we claim that the algorithm does not reconstruct relations between 1D-genetic marks and 3D-genome organization; instead, *TargetFinder* obtains information about 3D-interaction of each region from training dataset, and uses this information to predict interactions of regions belonging to validation dataset. To distinguish genomic regions and align regions in training and validation dataset, *TargetFinder* utilizes distribution of the 1D-genetic marks as region signatures. In this case, 1D-genetic marks within window between promoter and enhancer are the most informative predictors only because they allow to effectively discriminate genomic intervals, which does not have any biological relevance. Thus, *TargetFinder* would be able to “predict” EP interactions only if EP window is present in both training and validating samples, and fails to make prediction otherwise. It fits well with the fact that f1-score drops dramatically when EP pairs for training and validation datasets are sampled from different chromosomes (Table 1) or when pairs with shared promoter or enhancer are excluded from analysis (f1-score < 0.3).

It is important to note that other published algorithms also use similar design of training and validation datasets (see discussion). Thus, our results highlighted specific peculiarities of biological data, which should be considered in future to prevent overfitting issues and wrong efficiency estimation of machine-learning approaches focused on predictions of chromatin interactions.

### Enhancer-promoter interactions are quantitative rather than qualitative

As we found that *TargetFinder* cannot efficiently predict EP interactions, we aimed to improve this algorithm. We considered following enhancements.

First, we decided that epigenetic marks not only between, but also outside of, promoter and enhancer should be considered. This makes sense in light of recently discovered mechanisms, underlying spatial organization of chromatin. For example, according to the loop extrusion model ^14^, binding of CTCFs in converged orientation outside of, but close to, EP pair will result in increased looping between promoter and enhancer. Based on the loop extrusion model, we also introduced orientation of CTCF sites as a predictor.

Second, we override definition of enhancer-promoter *interaction*. According to the approach utilized previously, promoters and enhancer that occur within anchors of a Hi-C loop were considered as interacting, whereas all other EP pairs were considered as non-interacting. To benchmark this approach, we collected all interacting EP pairs for human monocytes based on SlideBase and GeneHancer databases (see methods for details). In addition to promotor-capture 3C data, these databases utilize information of co-expression of promoters and regulatory elements, their distance and other information to define interacting EP pairs. We choose human macrophage and monocytes data since information about these cell types was available in SlideBase and GeneHancer, and high-resolution Hi-C maps for these cells were recently published ^15^, which allowed comparison of Hi-C loops and interacting EP pairs. We confirmed that contact frequency between interacting EP pairs, as well as between loop anchors, is higher than average contact frequencies (Fig. 2 A,B). However, vast majority of interacting EP pairs do not overlap with loops, although they are often located within reasonable distance from loop (Fig. 2, C). Similar results were obtained for human macrophages (Supplementary Fig. 1)

**Fig 2.**
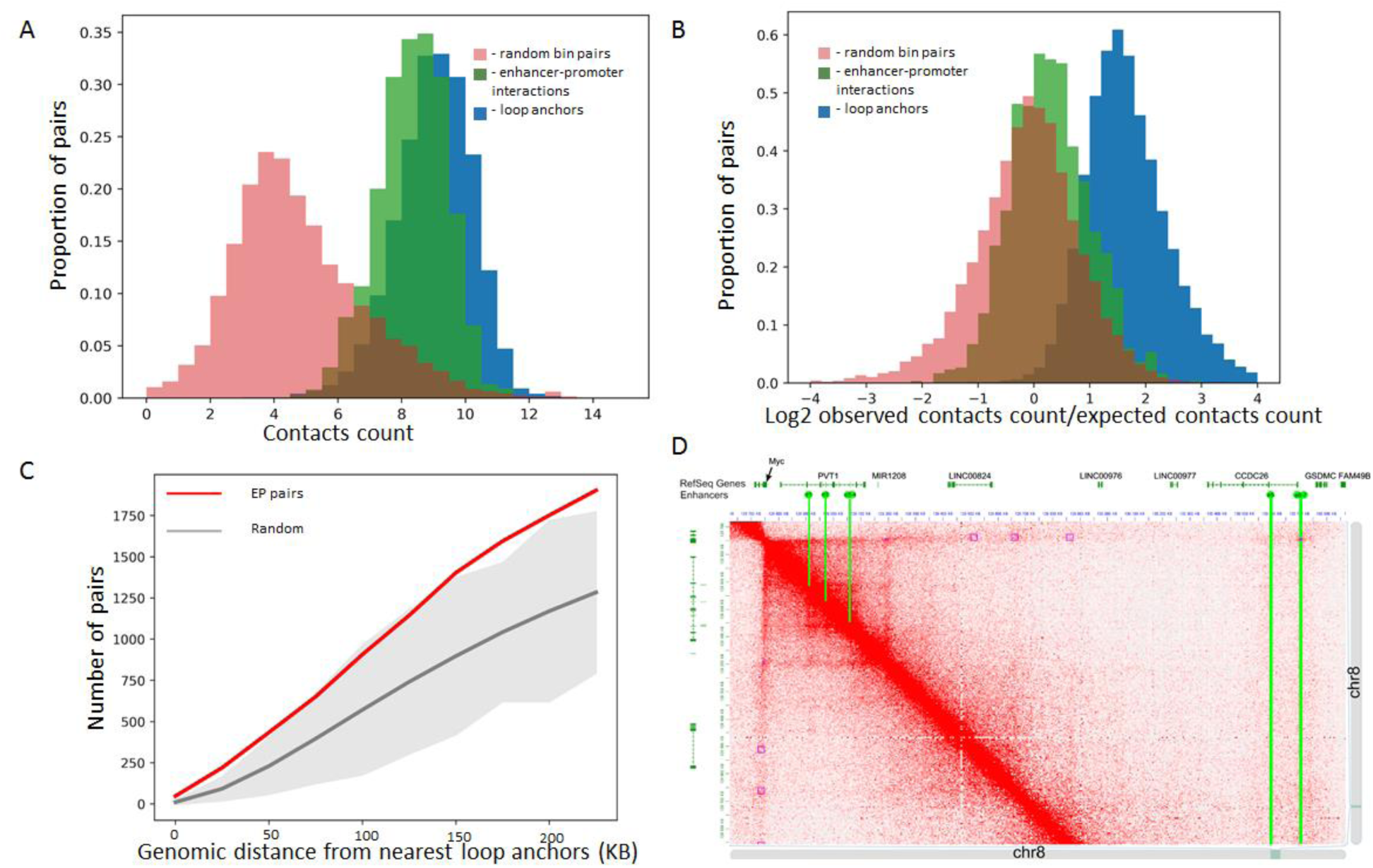
Hi-C loops do not provide complete information about interacting EP pairs. **A,B** Distribution of row (A) and distance-normalized (B) contact frequencies for interacting EP pairs and loop anchors in monocytes. **C.** Number of interacting EP pairs overlapping loops in monocyte data. Red line - number of EP pairs overlapping any loop or located within distance not more than *X* kb of loop anchors as function of *X.* Gray line and grey area represent average and 3 standard deviations of 100 samples, which were obtained randomly selecting pairs of loci which are characterized by same distance distribution as EP pairs.

The set of EP pairs described in SlideBase and GeneHancer is probably not complete, and these databases (partially) relay on the 3C information to infer enhancer-promoter connections. The gold-standard for identification of functional EP interactions is direct genetic screening. Such screenings are expensive and time-consuming, thus there is very limited number of experimentally validated enhancers, which prevents systematic genome-wide analysis of their relations to Hi-C loops. However, individual reports of genetically validated functional EP interactions support our general conclusion. For example, recent study ^16^ identified 7 distal enhancers of c-Myc gene, which form 2 clusters located 0.16 and 1.9 MB away from the c-Myc promoter, respectively. According to ^16^ all of these enhancers affect c-Myc expression in K562 cells proportionally to the number of contacts between enhancer and c-Myc promoter in this cell type. However, we found that only e6 and e7 enhancers overlap Hi-C loops (Fig. 2D). Moreover, out of 5 loops containing c-Myc promoter only one contains validated c-Myc enhancers. Altogether, this means that binary classification of EP interactions guided by location of Hi-C loop anchors may have poor predictive power. These observations are consistent with the recent model of enhancer-promoter communication ^17^, which suggests that loops and domains serve to decrease effective distance separating enhancers and promoters, but are not necessarily formed by EP pair itself.

Thus, we concluded that increased interaction frequency, rather than location within loop anchors, should be used as a judgment of EP interaction. As spatial interactions are quantitative, we aimed to design quantitative algorithm which predicts frequencies of spatial interactions between genomic loci in general, and EP interactions in particular.

### Quantitative prediction of EP interactions using machine learning approach

We used following biological information to predict EP and other genomic interactions: ChIP-seq profiles, describing chromatin binding of architectural proteins or histone modifications; RNA-seq profiles, describing gene expression levels; E1 values, classifying chromatin to active (A) and inactive (B) compartments; genomic distance, which is essential factor of 3-dimensional contacts. We restricted our algorithm to the prediction of mid-range contacts (<= 1.5 MB) since almost all EP interactions occur within this distance. To increase sample size and avoid overfitting, we included contacts of all loci, regardless of presence of promoters and enhancers, into the training set. We always performed training and validation on different chromosomes, and never used chromosome number or genomic coordinates of loci as predictors, to prevent overfitting.

Using recently generated Hi-C and genomic data on mouse hepatocytes, we compared several forms of predictor parametrization and performance of different machine learning algorithms by measuring as Pearson correlation, stratum adjusted correlation coefficient (SCC) ^18^, mean squared error (MSE), mean absolute error (MAE) and mean relative error (MRE) (see *Supplementary Note* for details). As a result, we developed *3DPredictor*, a machine-learning tool for computational prediction of chromatin interactions. Notably, *3DPredictor* provides accurate predictions using only CTCF ChiP-Seq data (including orientation of the occupied CTCF-sites), RNA-seq data and distance between loci as input, and only one chromosome out of 20 for training, as was evident from training and validation on mouse hepatocyte data (Pearson R=0.92-0.95, SCC=0.53-0.72, MSE=0.0017-0.0082, MAE=0.0010-0.0015, MRE=0.52-1.74; Fig. 3). These results can be further improved when multiple chromosomes were used for training (Fig. 3 B-D).

**Fig 3.**
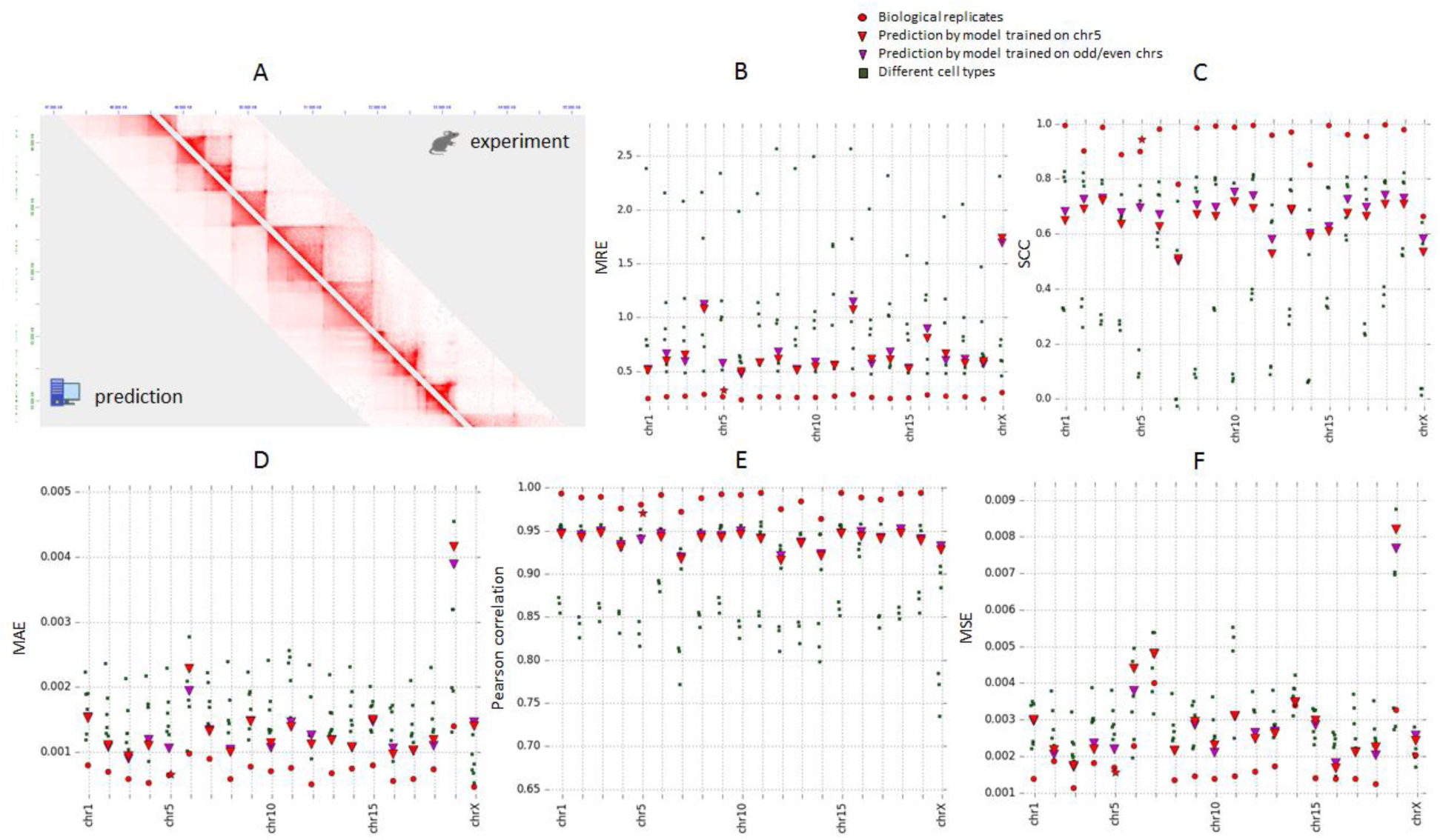
3DPredictor efficiently reconstructs spatial interactions based on CTCF-occupancy, expression and genomic distance. A. Representative region of mouse chromosome 2 showing predicted and experimentally derived Hi-C interactions in mouse hepatocytes. B-F. Various metrics of 3DPredictor accuracy for each chromosome of mouse hepatocytes. Circles represent comparison between two replicas; red squares shows comparisons between hepatocytes and other cell types. Red triangles display 3DPredictor results obtained using single chromosome 5 for training; data obtained when validating on the same chromosome marked with asterisks. Blue triangles show results of 3DPredictor trained on 10 chromosomes (results for even chromosomes obtained training on odd chromosomes and *vice versa*)

It is known that chromatin contacts are moderately similar between cell-types ^1,19^. To find whether our predictions are cell-type specific, we first compared chromatin architecture of different cell types using aforementioned measures. Importantly, in most of cases results obtained by 3DPredictor differ from real data less than cell types differ from each other (Fig. 3 B-D). For example, for 13 chromosomes 3DPredictor results, judged by mean average error, resemble experimental hepatocyte’ Hi-C data more closely than experimental data derived from other studied cell types. For 4 more chromosomes (chromosomes 1, 4, 6, 9 and 15) prediction errors were comparable with MAE obtained for different cell type, and on chromosomes 19 and X predictions were worse than transferring contact counts from other cell types. Similar results were obtained for MSE and MRE. According to the SCC, 3DPredictor performs approximately at the level of intercellular differences, whereas according to Pearson correlation predictions were almost always more similar to the hepatocyte’ data than other cell types.

We next compared Hi-C data of mouse hepatocytes and NPC and found that some genomic regions show apparently different 3D-organization in these cell types. In most cases, the differences were due presence of cell-type specific TADs, which borders coincide with cell-type specific CTCF sites, as was observed previously ^1,13^. We manually selected 14 representative regions characterized by strongly different chromatin organization, and run 3DPredictor for these regions using cell-type specific predictors. To measure how well cell-type specific TADs were captured by 3DPredictor, we calculated insulatory index around boundaries of cell-type specific TADs. As shown on Fig. 4, correlation of insulatory index derived from experimental and 3DPredictor data was very high for 11 out of 14 studied loci, which indicates that for cell-type specific regions our predictions are much more accurate than transfering contact counts from another cell type. An example of accurate prediction of cell-type specific TAD boundary in NPC and hepatocytes shown on Fig. 4 B and C.

**Figure 4.**
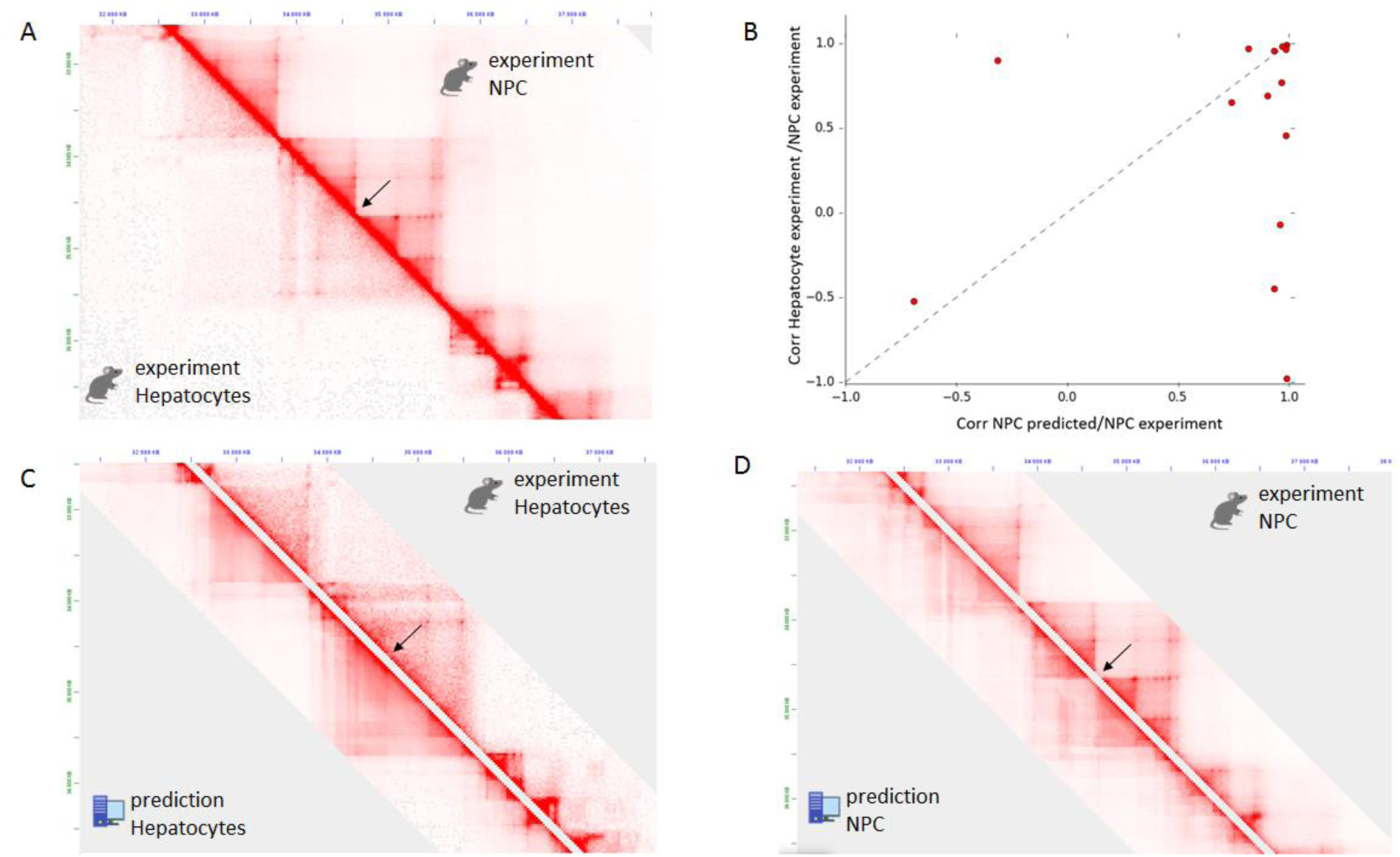
3DPredictor accurately reconstructs cell-type specific chromatin organization. A. Representative region on chromosome 3 showing different organization in mouse hepatocytes and NPC. Cell-type specific TAD boundary indicated by arrow. B. Insulatory index derived from either 3DPredictor or NPC data compared with hepatocyte’ insulation in 13 independent hepatocyte-specific regions. C and D. Comparison of 3DPredictor results with experimental hepatocyte (C) or NPC (D) data for the same region of chromosome 3 as on A.

Finally, we run 3DPredictor on human GM12878 data (Supplementary Fig. 2). According to all metrics except SCC, predictions fit experimentally-derived Hi-C interactions better than data from other cell types, even when using single chromosome for training. In the same time, transferring interaction frequencies from other cell types results in better SCC values comparing to model predictions with only one exception on chromosome 9, and, in general, SCC values obtained on human data were slightly lower than obtained on mouse data.

It is worse mentioning that EP interactions were predicted as good as other interactions. MRE of contact frequencies for interacting (according to SlideBase and GeneHancer databases) EP pairs was slightly lower than for all chromatin interactions predicted in monocytes, and MSE and MAE slightly lower (Fig. 5). In general, experimentally-derived contact frequencies of EP pairs in monocytes were highly correlated to predicted contact frequencies for corresponding loci in these cells (Fig. 5).

**Figure 5.**
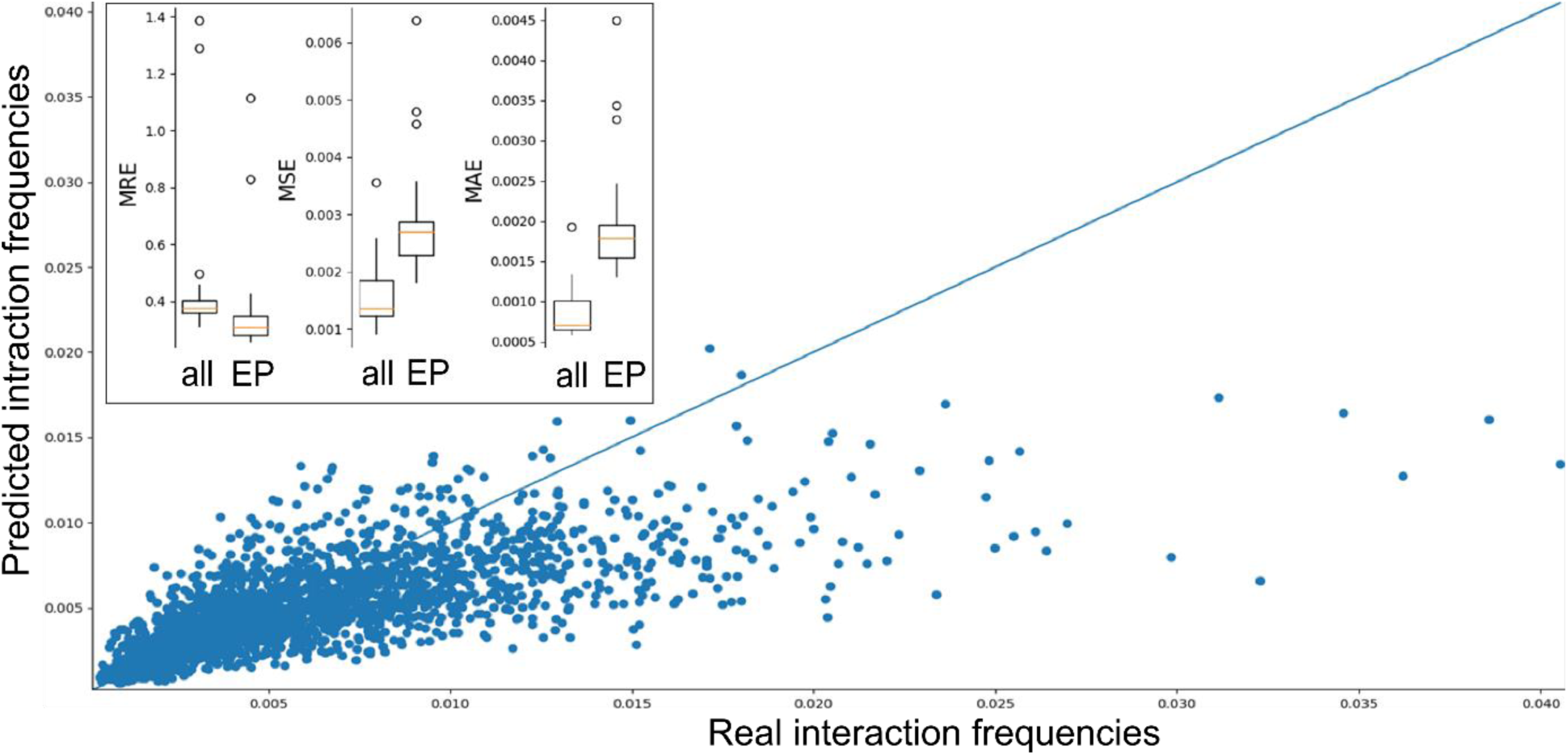
Accurate prediction of promoter-enhancer interaction frequencies. Contact frequencies for EP pairs defined as “interacting” in monocytes according to SlideBase and GeneHancer databases (X-axis) plotted against predicted contact frequencies (Y-axis). Estimates of prediction errors for all contacts and EP pairs shown within box in the top-left corner.

We next used 3DPredictor trained on mouse hepatocytes (single chromosome or half of genome) to predict contact frequencies in mouse NPC. Predictions of spatial interactions for cell type which was not used for training 3DPrediction appeared to be as good as when same cell type was used for training and validation (Fig. 6). From practical point of view, this indicates that our approach can be used to predict 3-dimensional genome organization, including EPI-contacts, in those cell types where 3C data is not available. From biological standpoint, these results show that principles of genome architecture are very similar in different cell types.

**Figure 6.**
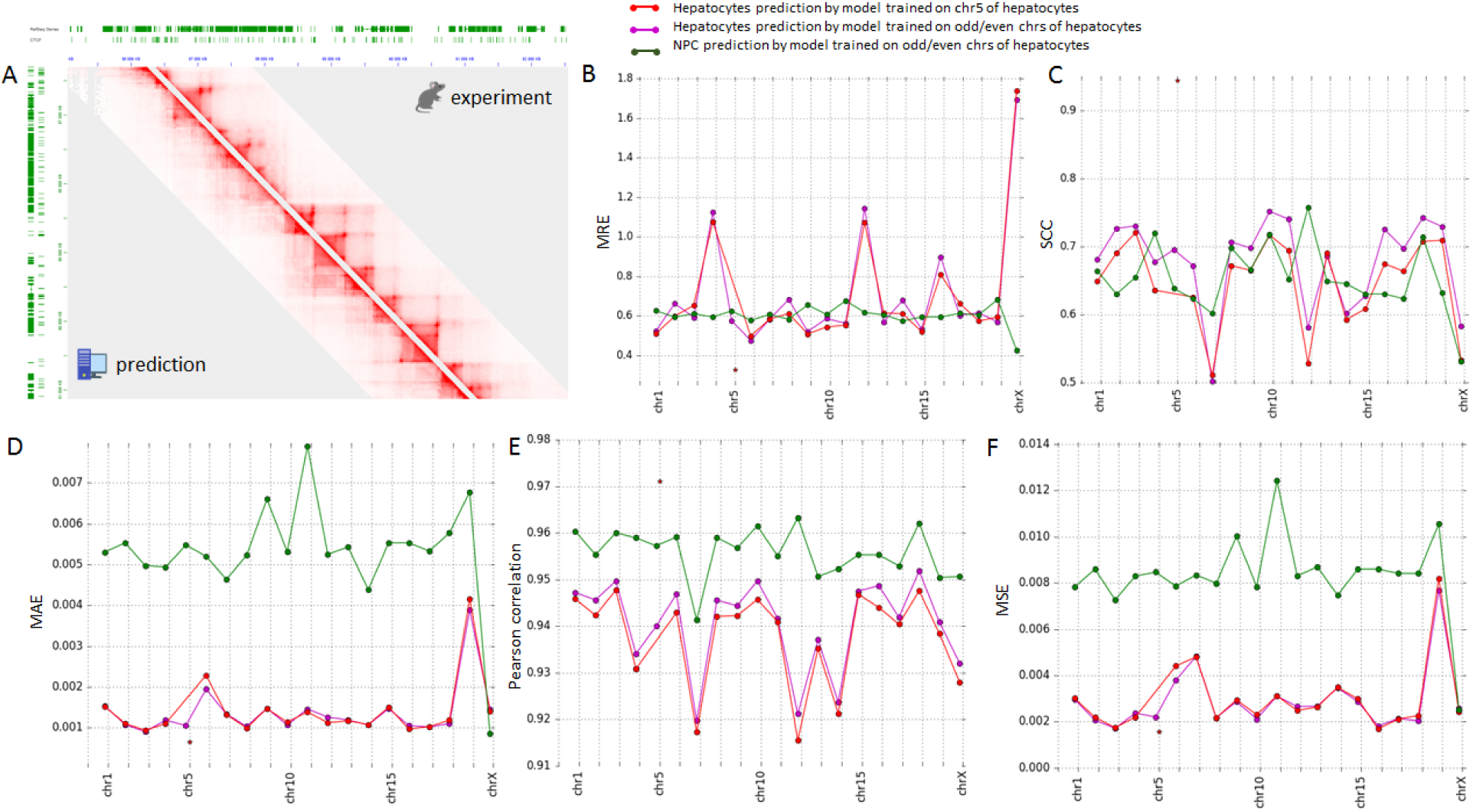
3DPredictor accurately reconstructs genome organization of novel cell type. A. Example of mouse NPC Hi-C contact map derived from experimental data (above diagonal) or obtained training 3DPredictor on hepatocyte’ contacts and providing epigenetic data relevant for NPC. B. MRE (B), SCC (C), MAE (D), Correlation (E) or MSE (F) estimate of 3DPredictor accuracy for training and validation on same (green and blue lines) or different (red line) cell types.

**Figure 7.**
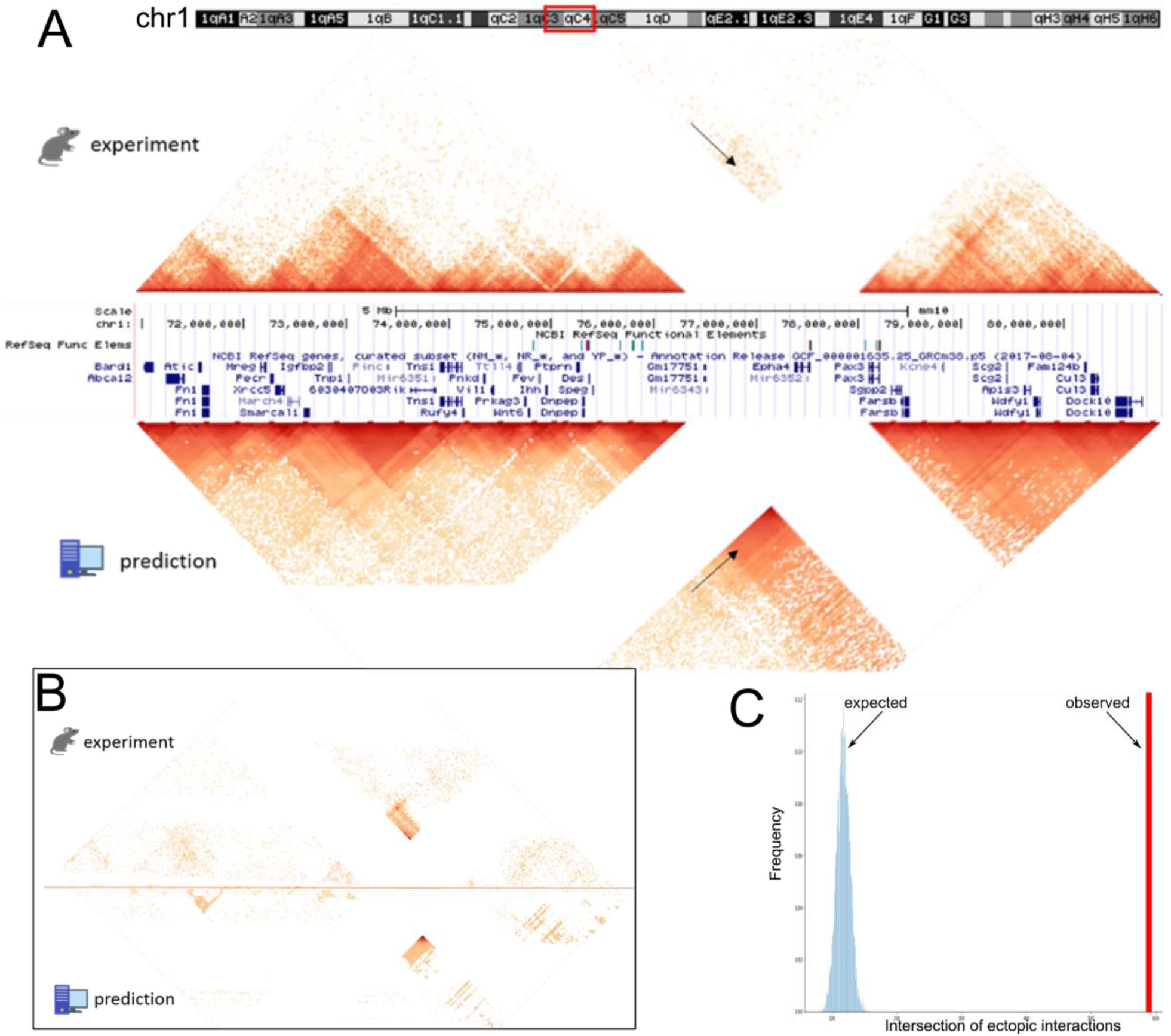
3DPredictor captures 3-dimensional organization of rearranged genomic regions. **A**. Contact map of mouse *DelB* locus, carrying homozygous deletion of ∼1.5 MB, with experimentally-measured contacts in the top, and 3DPredictor modeling results in the bottom. White region corresponds to contacts of deleted locus. Note ectopic interactions between Pax3 and Epha4 TADs (indicated by arrows). These ectopic interactions are even better visible on **B**, where the same region is plotted and only those interactions which differ between WT and *DelB* by more than 3 standard deviations are kept. On **A**, the color indicates contact counts, whereas on **B** the color indicates difference between WT and *DelB* data. **C** shows sizes of observed (red vertical bar) and expected (blue bars) overlaps between experimental and predicted ectopic interactions.

### Predicting effects of chromosomal rearrangements on 3-dimensional genome organization

One important application of enhancer targets prediction is understanding of EP reviewing after chromosomal rearrangements. There are several notable examples of pathological changes in EP contacts caused by deletions, inversions ^20^ or duplications ^21^. Recently, PRISMR ^22^ was developed to resolve chromatin structure of rearranged genome. Although impressively accurate, PRISMR requires Hi-C data to optimize chromatin model parameters, which limits its usage to cell types with available Hi-C data or genomic regions with 3-dimensional structure conserved across cell types. 3DPredictor lacks these limitations, as we have shown that it can predict chromatin packaging of cell-type specific regions and previously unstudied cell types.

We employed recently generated 5C data describing mouse EPHA4 rearrangements to find whether 3DPredictor can infer ectopic interactions in mutated genome. We re-analyzed 5C data generated from wild-type cells, as well as cells carrying homozygous deletion of ∼1.5 MB encompassing *Epha4* gene ^12^. This deletion (referred as *DelB* in ^22^) results in establishment of ectopic contacts between *Pax3* gene and *Epha4* enhancers cluster, which is associated with *Pax3* misexpression leading to brachydactyly.

We run 3DPredictor trained on mouse hepatocytes to infer 3-dimensional organization of the rearranged EPHA4 loci in hindlimb cells. Notably, although we did not use any *a priori* knowledge of 3-dimensional structure of wild-type EPHA4 locus in hindlimb cells, 3DPredictor results were very similar to experimental data (Fig. 5 A). We used method described in ^22^ to find ectopic interactions in the rearranged locus. Out of 1561 interactions inferred from the experimental data, 589 were captured by 3DPredictor, including majority of interactions between *Pax3* gene and *Epha4* enhancers (Fig. 5 A, B). The overlap between real and predicted ectopic interactions was large and strongly differ from randomized control (Fig. 5C, p-value < 5e10-6). This shows that our model successfully predicts ectopic interactions in the rearranged genome.

## Discussion

Machine-learning approaches are actively employed to capture complex epigenetic signatures underlying chromatin contacts. Although these methods can be extremely powerful, they do not provide mechanistic explanation of captured dependences, in contrast to the alternative approaches which utilize methods of physical polymer modeling. As we have shown here, biological data may have specific structure. For example, pairs of loci with overlapping window partially share epigenetic environment and often display similar 3-dimensional architecture. This means that these regions can not represent independent samples in training and validation datasets, otherwise correlations captured by machine-learning approaches may not reflect causations underlying genome architecture.

Notably, *TargetFinder*, which was used as a representative example here because it’s often cited as “golden standard” of tool for Enhancer-promoter interaction prediction, is not the sole example of research that does not take account of this peculiarity of biological data. For example, recently published *EP2vec* ^3^ utilizes same dataset as *TargetFinder* and constructs training and test samples in the same way. Another tool aimed to predict CTCF loops *CTCF-MP* ^12^ does not take into consideration nested loops when uses window features. Although both *EP2Vec* and *CTCF-MP* can generate predictions without window information, performance of such setup is lower: ∼10% of accuracy drops when *CTCF-MP* trained without DNAseI and ChIP-Seq window features and ∼2-4% of f1-score drops when EP2Vec trained without *TargetFinder*-derived window features.

In recent preprint describing a tool for HiC-data prediction *HiC-Reg* ^23^ authors also show that sharing genomic regions between training and validation datasets improve prediction scores. However, authors explain this observation as chromosome-specific biological mechanisms which cannot be modeled when data from predicting chromosome is omitted from training set. Whereas chromosome- and even region-specific mechanisms of DNA-packaging indeed exist ^24^, and we also shown that prediction is better when multiple chromosomes are used to train the model, it is also probable that better results of intrachromosomal cross-validation originate from existence of overlapping regions. One should note, that although pairs with overlapping left- or right-anchor are not shared between training and validation datasets in HiC-Reg (so-called easy samples), authors do not exclude regions which share part of window between interaction anchors.

We next raise the question of definition of promoter-enhancer *interaction*. Currently, most of studies use all 3C-interactions which differ statistically from distance-adjusted background as functional PE interactions ^5,25^. According to our results, functional interactions of promoters and enhancers do not fully overlap with Hi-C loops, and, probably, do not overlap completely with any other set of enriched interactions. Undoubtedly, spatial proximity is required for PE communication; however, it is not clear which spatial distance is necessary and sufficient to achieve functional interaction. For example, recent study of Shh-ZRS TAD showed that almost whole ∼900-kb intra-TAD region can be activated by ZRS enhancer, although pronounced looping was observed only between Shh promoter and ZRS enhancer ^26^. Removing Shh-ZRS TAD boundary reduces intra-TAD contact frequencies to the background level and disturbs Shh expression in the developing limbs; however, relocating enhancer closer to Shh promoter region restores expression pattern. These results indicate that background-level interaction within TAD might be sufficient to establish functional connections of promoter and enhancer. Moreover, recent paper reports that intra-TAD promoter regions often show significant level of interaction with TAD boundaries, and disruption of these interactions does not lead to changes of expression levels ^27^. To sum up, our view is that statistical increase of spatial contact frequencies, i.e. formation of loops, is important indicator of promoter-enhancer connectivity, but cannot be solely used to distinguish functional interactions. In accord with this, recent large-scale CRISPR assay of promoter-enhancer connections ^28^ suggested quantitative “contact-by-activity” model of PE interaction. In this model, enhancer impact is quantitative and proportional to both promoter-enhancer proximity and enhancer activity. Whereas letter can be estimated using DNAseI or ATAC-seq data available for many cell types, we developed 3DPrdeictor, a new approach to quantitatively predict spatial architecture of chromatin, including enhancer-promoter interactions.

Our analysis confirms that features describing window between interacting regions are essential for modeling interaction frequencies, in accord with observations described in *TargetFinder* paper ^5^. Importantly, we performed unbiased estimation of predictive power of our algorithm correctness, using correct training and validation datasets and various correlation- and distance-based metrics. We showed that using our approach one could predict spatial organization using very limited amount of input data, such as CTCF ChIP-Seq tracks and genome-wide expression obtained by RNA-Seq analysis. Importantly, we benchmarked our predictions against transferring contact frequencies from different cell types. Although modeling outperforms this approach for most of cell types and chromosomes, it is not yet accurate enough to achieve similarity level of biological replicates.

It is essential that our model not only predicts chromatin interactions in normal genome, but also can capture ectopic interactions which arise as a result of chromosomal rearrangements. Currently, both experimental data describing 3D-genome alterations for known rearrangements and tools modeling spatial landscape of novel variants are limited. In the same time, others ^20,21,29,30^ and we ^31^ have recently reported novel variations with unexpected pathological phenotype, which might be explained, at least partially, by changes of chromatin organization ^32,33^. Future development and validation of models predicting chromatin contacts in rearranged genome is essential for better understanding of biomedical consequences of these rearrangements. Moreover, integrating chromatin interactions, derived from 3DPredictor, with enhancer activity information using the “activity-by-contact” model may allow precise estimation of transcriptional changes caused by structural variations.

## Materials and Methods

### Hi-C data processing

Hi-C data for mouse hepatocytes (GSE95116) was downloaded from NCBI and processed using JuicerTools ^34^. Resulting .hic files are deposed at genedev hic-files server (http://genedev.bionet.nsc.ru/site/hic_out/) under accession “Hepat”. Hi-C data for mouse ES cells and neural precursors ^13^, and human GM12878 ^1^, K562 ^1^, macrophages ^15^ and monocytes ^15^ were available at AidenLab hic-files server via JuiceBox and JuicerTools. All datasets were KR-normalized. For each Hi-C dataset, contacts were at 25-kb resolution using JuicerTools *dump* command. To be able to perform comparisons between cell types, we normalize datasets dividing each contact by normalization coefficient *Coef,* which reflects average bin coverage:

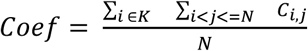, where *C*_*i,i*_-contacts between i-th and j-th bins, K - number of bins on chromosome 1, *N* - number of bins in genome. To speed up computations we only use bins of chromosome 1 while computing *Coef*, although this should not affect results as we use KR-normalized matrices where coverage of all bins are roughly equal.

Loops were called by JuicerTools *hiccups* command with default parameters using heatmaps at 25-kb resolution. K562 loops presented on Fig. 2 are from ^1^.

First eigenvector (E1) values of Hi-C matrixes were obtained using JuicerTools *eigenvector* module.

5C data describing 3-dimensional organization of wild-time and mutated mouse Epha4 locus in distal limb buds were downloaded from GEO: GSE92291. Data was processed by HiCPro ^35^ pipeline using mm10 genome.

The relative error (RE) of Hi-C contact counts shown at Supplementary Fig. 3 was accounted based on binomial distribution: 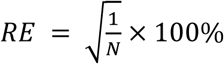, where *N* is a number Hi-C reads between contacted loci. The average RE and standard deviation of REs were independently calculated for each genomic distance.

To estimate correlation of contact counts on different resolutions for Supplementary Fig. 4, we used data for chromosome 10 of GM12878 cell line. We randomly choose 1000 loci pairs and calculated Pearson’s correlation between KR-normalized contact frequencies on different resolutions. We aggregated calculations of 100 independent samplings by averaging to obtain final results.

### Definition of promoters and enhancers

For macrophage and monocyte enhancers were defined using SlideBase (http://slidebase.binf.ku.dk) database. This database is supported by the FANTOM5 consortium and represents a map of human regulatory elements of each cellular state. It contains levels of enhancer expression based on CAGE sequencing RNA isolated from every major human organ, over 200 cancer cell lines, 30 time courses of cellular differentiation, mouse developmental time courses and over 200 primary cell types. Thereby, an enhancer can be specific to a set of primary cells and organs (tissue samples) or can be broadly (or ubiquitously) expressed. We took into account enhancers displayed more than 25% emitting for target cell line.

Using GeneHancer database (https://www.genecards.org), we define gene promoters regulated by given enhancer. GeneHancer is a database of genome-wide enhancer-to-gene and promoter-to-gene associations, embedded in GeneCards. Regulatory elements were mined from the following sources:

1. The ENCODE project Z-Lab Enhancer-like regions
2. Ensembl regulatory build
3. FANTOM5 atlas of active enhancers
4. VISTA Enhancer Browser enhancers validated by transgenic mouse assays.
5. dbSUPER super-enhancers.
6. EPDnew promoters.
7. UCNEbase ultra-conserved noncoding elements.

GeneHancer associations and likelihood-based scores were generated using information that helps link regulatory elements to genes:

1. eQTLs (expression quantitative trait loci) from GTEx
2. Capture Hi-C promoter-enhancer long range interactions
3. Expression correlations between eRNAs and candidate target genes from FANTOM5
4. Cross-tissue expression correlations between a transcription factor interacting with an enhancer and a candidate target gene
5. GeneHancer-gene distance-based associations, scored utilizing inferred distance distributions. Associations include several approaches: (a) Nearest neighbors, where each GeneHancer is associated with its two proximal genes; (b) Overlaps with the gene territory (Intragenic); (c) Proximity to the gene TSS (<2kb). TSS proximity scores are boosted to elevate GeneHancer associations in the vicinity of the gene TSS.

The “true” interacting EP pairs of monocytes and macrophages were calculated by combining list of cell type specific list of enhancer from SlideBase and list of enhancer-gene associations from GeneHancer.

When using *TargetFinder* pipeline on human data, we used authors definition of active promoters and enhancers, and obtained coordinates from https://github.com/shwhalen/targetfinder. For mouse data we define interact pairs as promoters and enhancers located within the same loop anchor. We used TSS (Transcription Start Sites) downloaded from UCSC as promoters and active enhancers from ^36^ as enhancers.

### ChIP-Seq data processing

All ChIP-seq data for human GM12878 and K562 cell lines were downloaded from https://github.com/shwhalen/targetfinder. ChIP-seq data for mouse hepatocytes (NCBI SRX2578761-SRX2578762), mouse NPC (NCBI SRX2636706-SRX2636707) and human monocytes (ENCODE ENCSR000ATN) were downloaded from NCBI or ENCODE and processed using aquas pipeline (https://github.com/kundajelab/chipseq_pipeline).

### RNA-Seq data processing

RNA-seq data for human GM12878 (ENCODE <u>ENCFF212CQQ</u>) and human K562 (ENCODE <u>ENCFF026BMH</u>) cell lines were downloaded from ENCODE. RNA-seq data for mouse hepatocytes (NCBI GSE95111) and mouse NPC (NCBI GSM2533845) were downloaded from NCBI. Data for human monocytes (NCBI SRX2785183) were downloaded from NCBI and processed using standard protocols with HISAT2 and StringTie.

### Prediction of 3-dimensional interaction frequencies

For training purposes, all data was split into non-overlapping genomic intervals. Usually, we use one or several chromosomes for training, and other chromosomes for validation. To perform prediction genome-wide, we first used odd chromosomes for training and made predictions for contacts on even chromosomes, and then used even chromosomes for training and predicted contacts on odd chromosomes. If other is not mentioned, we used only CTCF and RNA-seq data for predictions. For all results except those described in Supplementary Note we used Gradient Boosting with parameters n_estimators=100, max_depth=9, subsample=0.7. Predictors parametrization and other details are explained in details in the Supplementary Note.

### Estimating predictions efficiency

We used several metrics to choose the best model. Pearson correlation is the most common metric, but unfortunately it is too sensitive to the dependence of the average contact amount on the current distance in the Hi-C matrices. We used the SCC metric ^18^ in order to more contrast highlighting other features of the matrix, except the dependence on distance. This is a special metric for comparing Hi-C matrices, based on Pearson correlation. To reduce the effect of random noise, we smoothed Hi-C matrices before calculating SCC, as was suggested by ^18^. All comparisons were carried out with the same noise smoothing parameter *h = 2* (see Supplementary Note and Supplementary Table 4 for justification of *h* value). In addition, to evaluate the model’s quality, we used other metrics such as MSE, MAE and average relative error.

### Modeling chromosomal rearrangements

To model chromosomal we used model trained on mouse hepatocyte cells. To generate predictors, we obtained CTCF (NCBI SRX1975285-SRX1975286) and RNA-seq (NCBI SRX1975216-SRX1975217) data from wild-type mouse hindlimb E11.5 cells. Next, we deleted all CTCF peaks and genes from the region [mm10]: **chr1**:76392403-78064264, which corresponds to the deletion coordinates described in ^22^. Resulting set of predictors was used to model all chromatin contacts within the region [mm10]: **chr1**:70950000-81000000. To compare contact frequencies predicted by the model with experimental data, we defined ectopic interactions as described in ^22^. We first generated normalized difference matrix between mutated and WT matrices. For this, we multiply the mutant matrix by a coefficient that equalizes the reads count equivalent of the regions that are not involved in the mutation, then we subtracted from the mutated matrix the WT matrix. We normalized the difference matrix by dividing each sub-diagonal by the average WT reads count at its corresponding pairwise genomic distance. Next. we find ectopic interactions for each sub-diagonal. Specifically we filter out the points above 96th percentile and calculate standard deviation of remaining values. All points which differ from zero by more than 3 standard deviations were considered ectopic.

### Code availability

3DPredictor source code (https://github.com/labdevgen/3DPredictor) and Jupiter Notebook with code used to reproduce TargetFinder results (https://github.com/labdevgen/targetFinderTests) are both freely available on github.

## Authors contribution

V.F. proposed the study. P.B., D.F. and V.F. benchmarked targetFinder algorithm. V.S. and P.B. wrote main parts of 3DPredictor code. E.M. implemented metrics of algorithm performance. V.S., P.B., E.M. and M.N. developed different predictor parameterizations and compared training parameters. E.M. implemented metrics of 3DPredictor performance. M.N. prepared training and validation data and analyzed EP interactions and loops in monocytes and macrophages. P.B. performed Epha4 locus modeling. All authors contributed to manuscript preparation.

## Acknowledgments

This work was supported by Russian Foundation For Basic Research (RFBR) grant #18-29-13021 and Russian Science Foundation grant #17-74-10143. All computations were performed with support from the Computational Cluster of the Novosibirsk State University and Computational Nodes of the Institute of Cytology and Genetics (Budget Project #0324-2019-0041). We acknowledge scientific discussions with Dr. N. Battulin.

## Supplementary material

### Supplementary Note. Effects of predictors parametrization and training parameters on prediction accuracy

We aimed to model interaction frequency for a pair of loci, L1 and L2, separated by genomic region (window) W12. We wish to make predictor parametrization so that it can describe chromatin structure of L1, L2, region W12 between them, and genomic neighborhoods surrounding L1 and L2. We note that promoters and enhancers are small features, often ∼1 kb in size. However, almost no Hi-C maps are currently available at this resolution, and for those that are available (such as, for example human GM12878 map) there is significant noise level at 1-kb resolution, which may affect model fitting (see estimations of noise level at Supplementary Fig. 3). For example, relative error of contact counts calculated for GM12878 data at 25-kb resolution is ∼10-20%, whereas for 5-kb resolution it is much higher (30-40% for distances below 400 kb, 40-70% for distances in diapason 0.4-1.5 Mb), and for 1-kb resolution average error raises to 80-90% for distances above 100 Kb (Supplementary Fig. 3). In the same time, according to the GM12878 Hi-C data, contacts at 25-kb correlate with the frequency of Enhancer-promoter interactions defined at 1-kb resolution reasonably well (25kb vs 5kb: R=0.75; 25kb vs 1kb: R=0.45; 5kb vs 1kb: R=0.73; Supplementary Fig. 4). Thus, we decided to predict contact frequencies at 25-kb resolution.

We initially performed following parametrization: for each pair of loci, L1 and L2, we defined a large genomic interval *I* of size 1.5 MB so that region L1-L2 would be located in the center of this interval. For each 25-kb bin within the interval *I* we compute average values of all available predictors and utilized obtained values as input. At this point we used following predictors a.) genomic distance between L1 and L2, b.) CTCF ChIP-seq signal value for each 25-kb bin and c.) the first eigenvector value (E1 value) derived from Hi-C matrix for each bin. One should note that E1 values were introduced as predictors only at this initial step as simple numeric representation of epigenetic state of each bin. In final models we do not use E1 values, since they cannot be obtained without Hi-C matrix and because by definition of E1 values their multiplication should give a good estimation of interaction frequencies.

Using this parametrization, we generate 121 values describing epigenetic state and CTCF distribution for each of 60 bins in the 1.5MB neighborhood of interacting loci. Although simple multiplication of E1 values should be informative for deciphering contact frequency, this parametrization showed poor performance of Linear regression, Gradient Boosting regression and Random Forest Regressor (Supplementary Fig. 5). Thus, we proposed another predictors parametrization.

For CTCF dataset, we generated following sets of features. Predictors describing genomic environment outside of the loci L1 and L2: i) distance to L1 and signal values of 4 peaks located on the left side of the L1 region (8 float values); ii) distance to L2 and signal values of 4 peaks located on the right side of L2 region (8 float values). Predictor describing genomic environment between loci L1 and L2: iii.) sum of signal values of all peaks within window W12 between interacting loci L1 and L2 (single float value). Predictors, describing loci L1 and L2: iv.) sum of all signals within 25-kb neighborhood of loci L1 or L2 (two float values). For E1 data, we designed predictors similar to iii.) and iv.), but instead of summing E1 values of bins we averaged values. Lastly, we used distance between L1 and L2 as a predictor (single integer value).

As loops are formed predominantly between CTCF-sites in convergent orientation, we decided to additionally parameterize CTCF sites orientation. For this aim, we introduced following predictors: 1.) for each CTCF peak described by predictors i) and ii) we added GimmeMotifs ^37^ motif score of forward and reverse orientation. For those peaks where CTCF motif was not found we set the score to 0; 2.) we described up to 8 CTCF peaks located between L1 and L2 in the same way as in predictors i) and ii) (for each peak we described distance to L1 or L2, signal value and forward/reverse motif score). Finally, we calculated number of convergently oriented CTCF blocks within the L1-L2 window. For this aim, we collapsed all consequentially located CTCF sites with the same orientation into one mega-site, and counted number of pairs of mega-sites with convergent orientation within the window W12.

Finally, we also decided to use RNA-seq data for predictions. RNA-seq data was parameterized in the same way as ChIP-seq data, with FPKM values used instead of peak intensities.

Using RNA-seq, CTCF binding and orientation and E1-values in as described above visibly improve Gradient Boosting model (Supplementary fig. 6). The model was able to predict TAD-like structures located at the similar positions as in experimental data. We next discarded E1-values, and found that this does not impair predictions (Supplementary fig. 6).

We than downloaded from ENCODE all data available for GM12878 cells, including ChiP-seq, FAIRE-sew, DNAse-sensitivity, methylation and etc. Although the model trained on a full set of 1909 predictors gave the best results for human GM12878 (Supplementary Table 1), feature importance analysis indicates that CTCF and RNA-seq are the most important predictors. Indeed, limiting input information to CTCF and RNA-seq data only results in only slight decrease of prediction rate. Similarly, the model trained on the full set of 123 predictors available for mouse hepatocytes performs better than model based on CTCF and RNA-seq only, however the difference in performance was reasonably small.

We also tested effect of training sample size on prediction accuracy (Supplementary Table 2). While larger training sets results in better performance, this dependence plateaued at ∼300 000 contacts, without any further increase of prediction accuracy when increasing sample size above this point. However, we note that using contacts sampled from different chromosomes performed better at this sample size (see main results).

We next compared Gradient Boosting with another ensemble learning method – Random Forest (Supplementary Table 3). Interestingly, Random Forest performance was significantly lower than of Gradient Boosting. As for different implementations of Gradient Boosting, we compared sklearn and xgboost implementations and found almost no difference (Supplementary Table 3). We also tweaked of xgboost hyperparameters, such as number of estimators, maximum depth of a tree and subsampling ratio, to achieve optimal performance.

Stratum adjusted correlation coefficient (SCC) was the main metric for the evaluation of algorithm performance. We used different smoothing factors for the matrix to improve accuracy of SCC. As evident from Supplementary Table 4, increasing smoothing factor *h* results in increase of SCC with, plateau around h=2. Thus, we chose smoothing factor equal to two and used this values of smoothing factor for all comparisons.

### Supplementary Figures

**Supplementary Figure 1.**
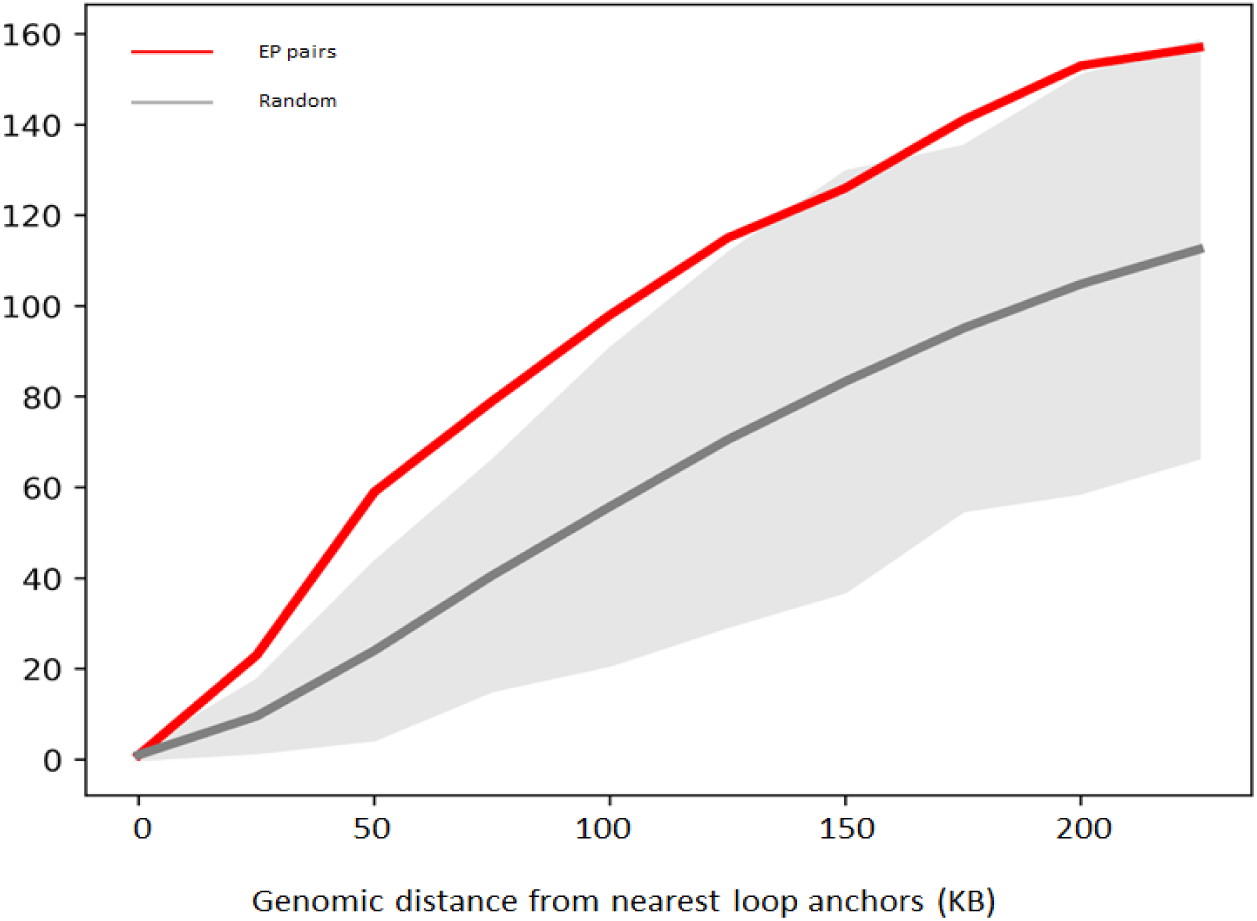
Promoter-enhancer interactions do not overlap Hi-C loops. Data for human macrophages presented as on Fig. 2C

**Supplementary Figure 2.**
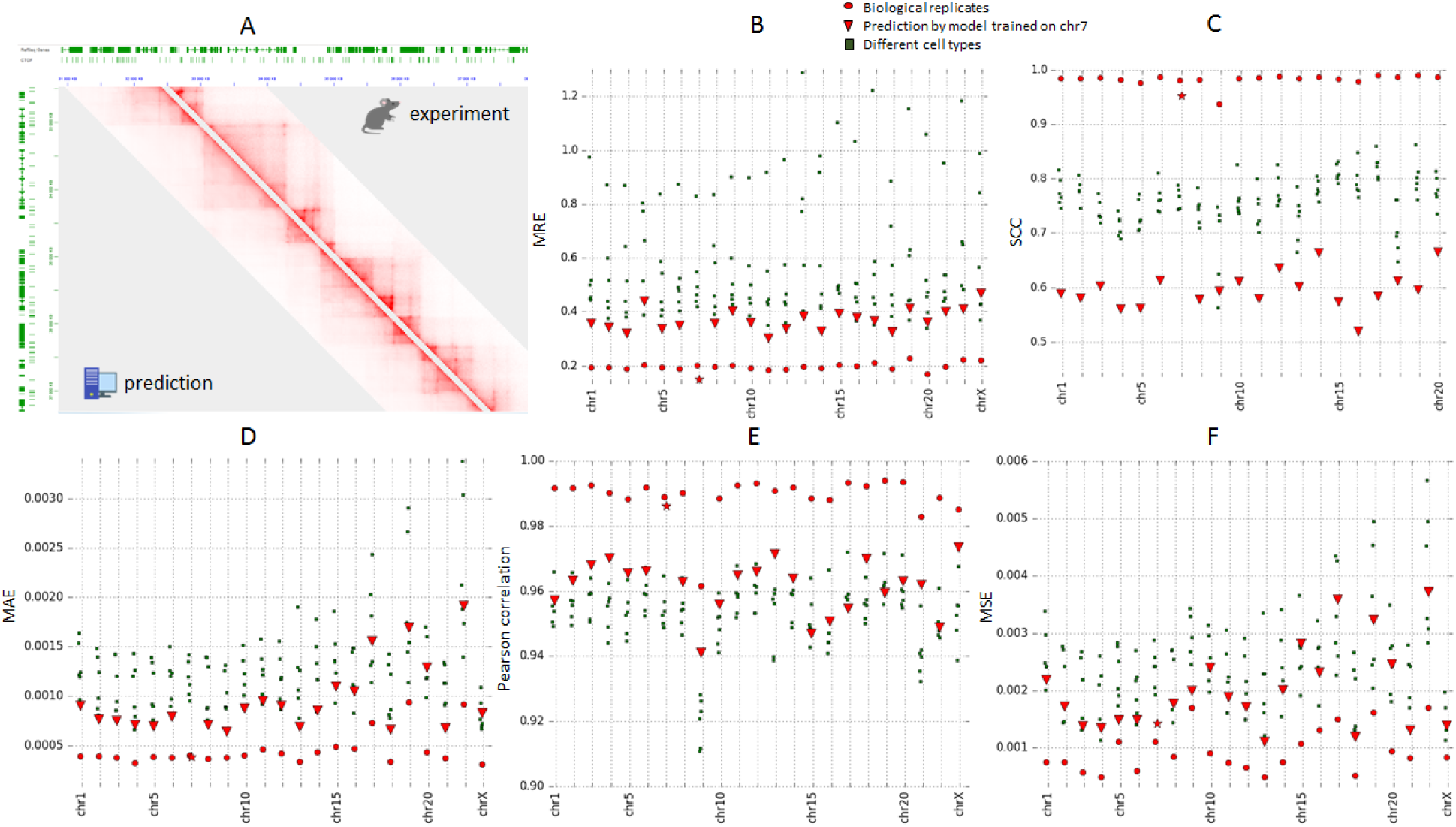
3DPredictor performance on human GM12878 data. Results are shown as on Fig. 3

**Supplementary Figure 3.**
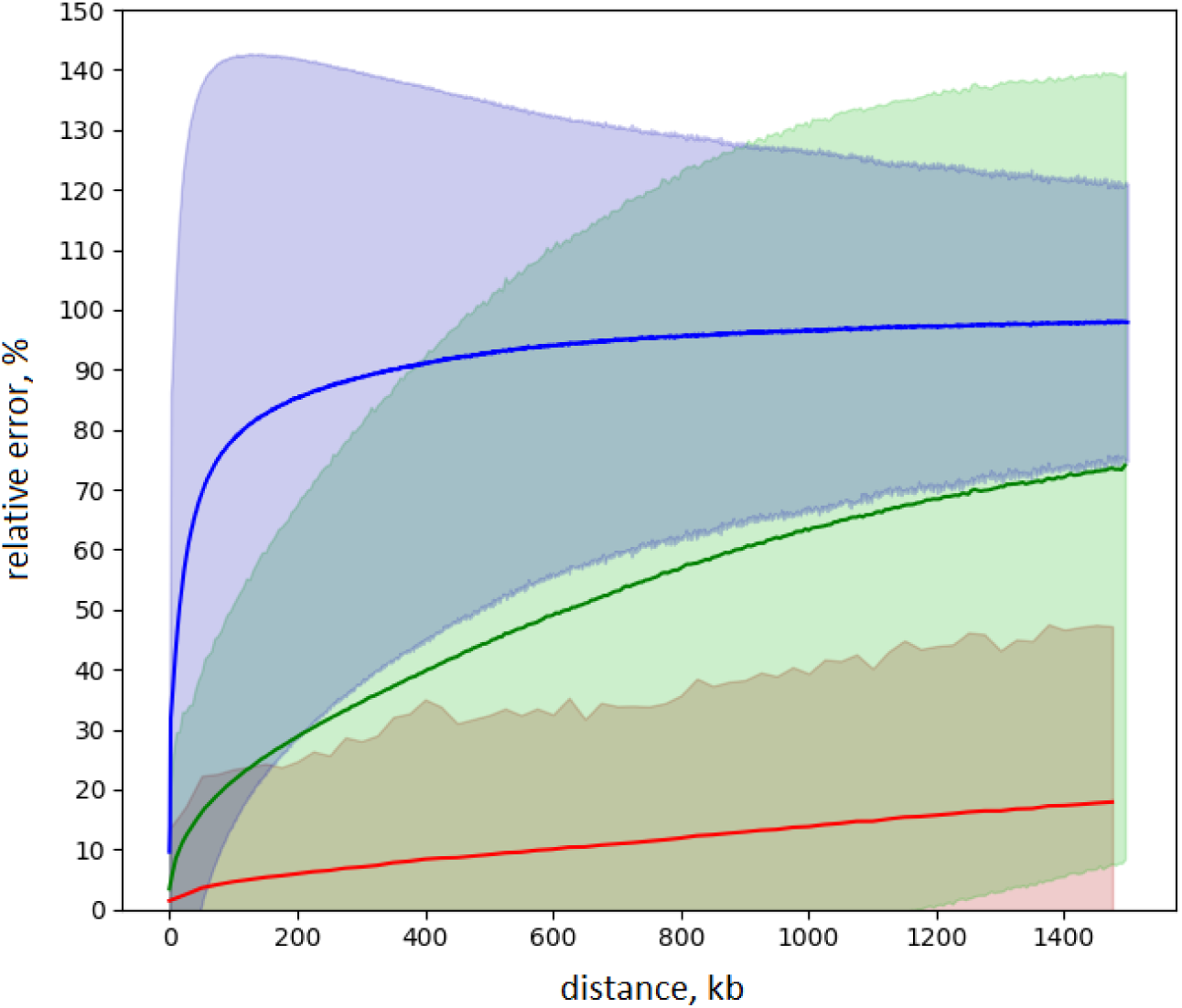
Dependence of Hi-C contact count error from genomic distance and resolution of analysis. The graph shows average (line) and 3 standard deviations (colored area) of the observational errors of Hi-C contact counts for contacts sampled at different distances (X-axis) and different resolutions (green - 25kb, red - 5kb, blue - 1kb). The data is presented for human GM12878 cells.

**Supplementary Figure 4.**
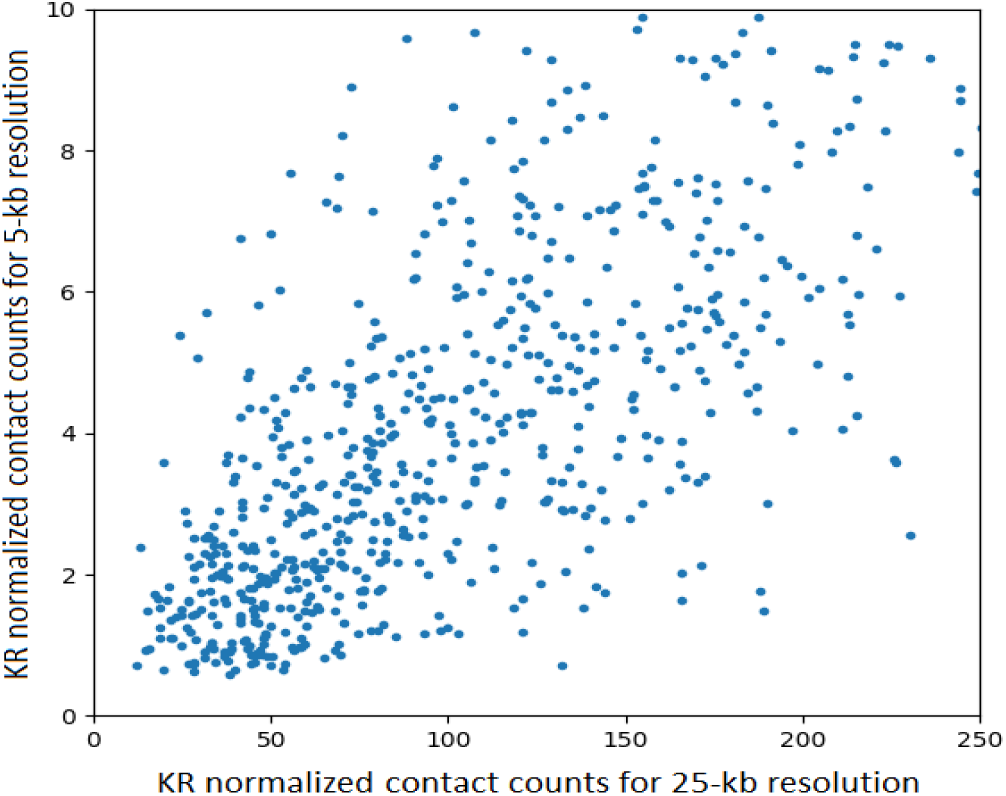
Dependence between KR normalized contact counts on different resolutions

**Supplementary Figure 5.**
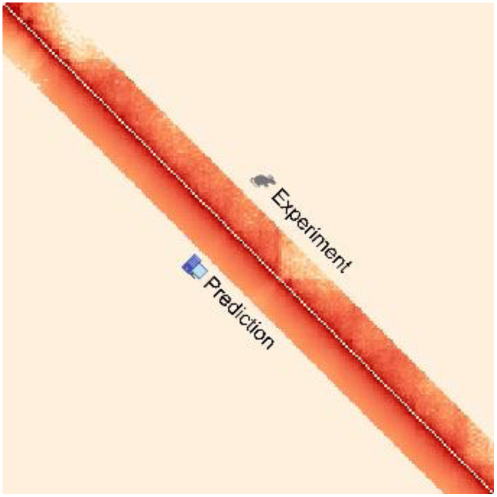
Poor prediction efficiency using initial parametrization describing epigenetic state and CTCF distribution for each of 60 bins in the 1.5 MB neighborhood of interacting loci.

**Supplementary Figure 6.**
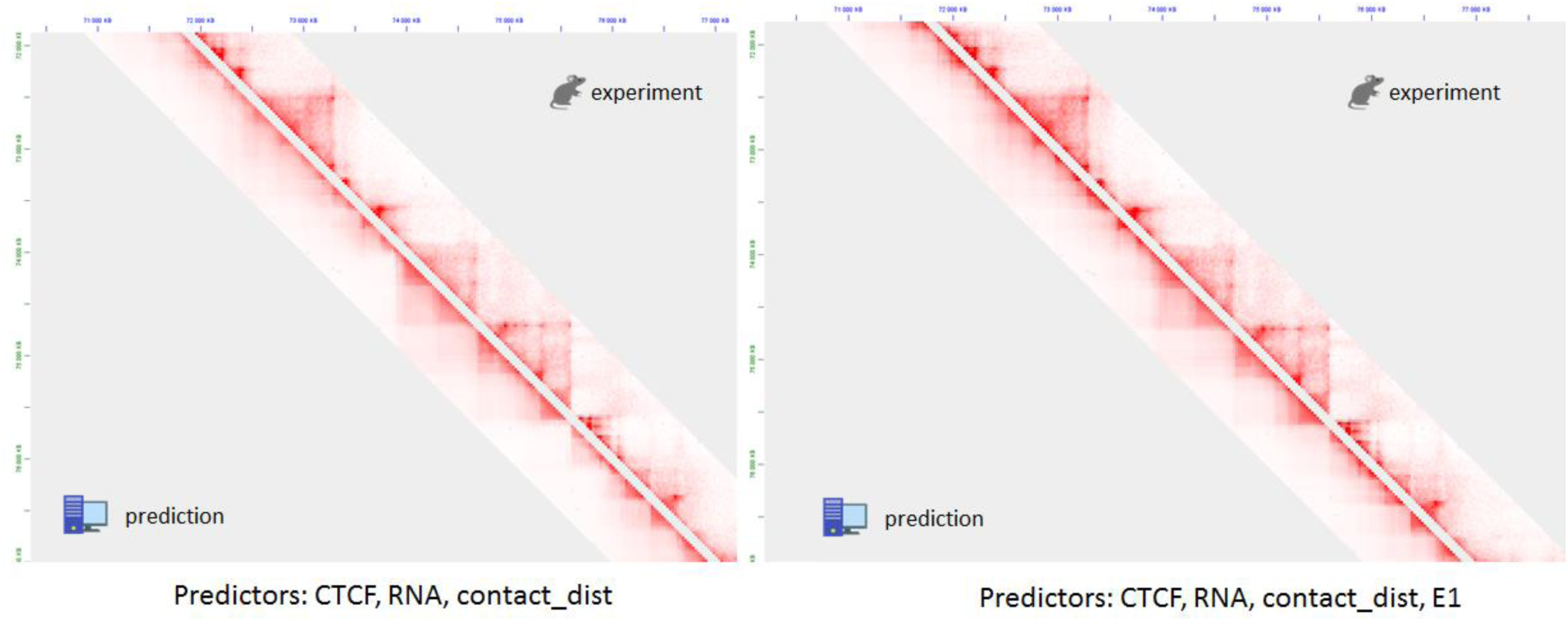
Representative region on chromosome 2 shows efficiency of 3DPredictor trained and validated on mouse hepatocyte’ data with (right panel) or without (left panel) using E1-values

**Supplementary Table 1.** 3DPredictor efficiency on GM12878 data depends on input epigenetic information.

| train size | Train regions | predictors | Validate regions | Pearson correlation | SCC | MSE | MAE | MRE |
| --- | --- | --- | --- | --- | --- | --- | --- | --- |
| 1 850 471 | Chr2 | all | Chr3 | 0,9707 | 0,6403 | 0,0013 | 0,0007 | 0,3288 |
| 1 850 471 | Chr2 | CTCF, CTCF_orientation, RNA-seq, contact_distance | Chr3 | 0,9685 | 0,6068 | 0,0013 | 0,0007 | 0,3387 |
| 1 850 471 | Chr2 | NFYB | Chr3 | 0,7912 | 0,0139 | 0,0041 | 0,0025 | 1,3259 |
| 1 850 471 | Chr2 | CTCF, CTCF_orientation, contact_distance | Chr3 | 0,9673 | 0,5769 | 0,0014 | 0,0008 | 0,3488 |
| 1 850 471 | Chr2 | FAIRE-seq | Chr3 | 0,8932 | 0,1903 | 0,0024 | 0,0014 | 0,5813 |
| 1 850 471 | Chr2 | DNAse and methylation | Chr3 | 0,8778 | 0,2362 | 0,0025 | 0,0015 | 0,7037 |

**Supplementary Table 2.** Effect of training sample on 3DPredictor efficiency.

| train size<br>(number of<br>contacts) | Train regions | predictors | Validate<br>regions | Pearson<br>correlation | SCC | MSE | MAE | MRE |
| --- | --- | --- | --- | --- | --- | --- | --- | --- |
| 25 000 | Chrms<br>1,5,9,13 | CTCF, CTCF_orientation, RNA-seq,<br>contact_distance | chr2 | 0,9389 | 0,7235 | 0,0022 | 0,0011 | 0,7051 |
| 25 000 | chr10 | CTCF, CTCF_orientation, RNA-seq,<br>contact_distance | chr2 | 0,9378 | 0,6610 | 0,0022 | 0,0012 | 0,7127 |
| 250 000 | Chrms<br>1,5,9,13 | CTCF, CTCF_orientation, RNA-seq,<br>contact_distance | chr2 | 0,9451 | 0,7359 | 0,0021 | 0,0011 | 0,6746 |
| 250 000 | chr10 | CTCF, CTCF_orientation, RNA-seq,<br>contact_distance | chr2 | 0,9257 | 0,7148 | 0,0000 | 0,0012 |  |
| 500 000 | Chrms<br>1,5,9,13 | CTCF, CTCF_orientation, RNA-seq,<br>contact_distance | chr2 | 0,9459 | 0,7356 | 0,0020 | 0,0011 | 0,6702 |
| 3 802 764 | Chrms<br>1,5,9,13 | CTCF, CTCF_orientation, RNA-seq,<br>contact_distance | chr2 | 0,9465 | 0,7345 | 0,0020 | 0,0011 | 0,6710 |
| 4 758 074 | All chrms<br>except 2, 19,<br>X | CTCF, CTCF_orientation, RNA-seq,<br>contact_distance | chr2 | 0,9468 | 0,7406 | 0,0021 | 0,0011 | 0,5916 |
| 917 632 | chr5 | CTCF, CTCF_orientation, RNA-seq,<br>contact_distance | chr2 | 0,9423 | 0,6490 | 0,0030 | 0,0015 | 0,5081 |
| 806 553 | chr10 | CTCF, CTCF_orientation, RNA-seq,<br>contact_distance | chr2 | 0,9418 | 0,6721 | 0,0021 | 0,0012 | 0,7127 |

**Supplementary Table 3.**
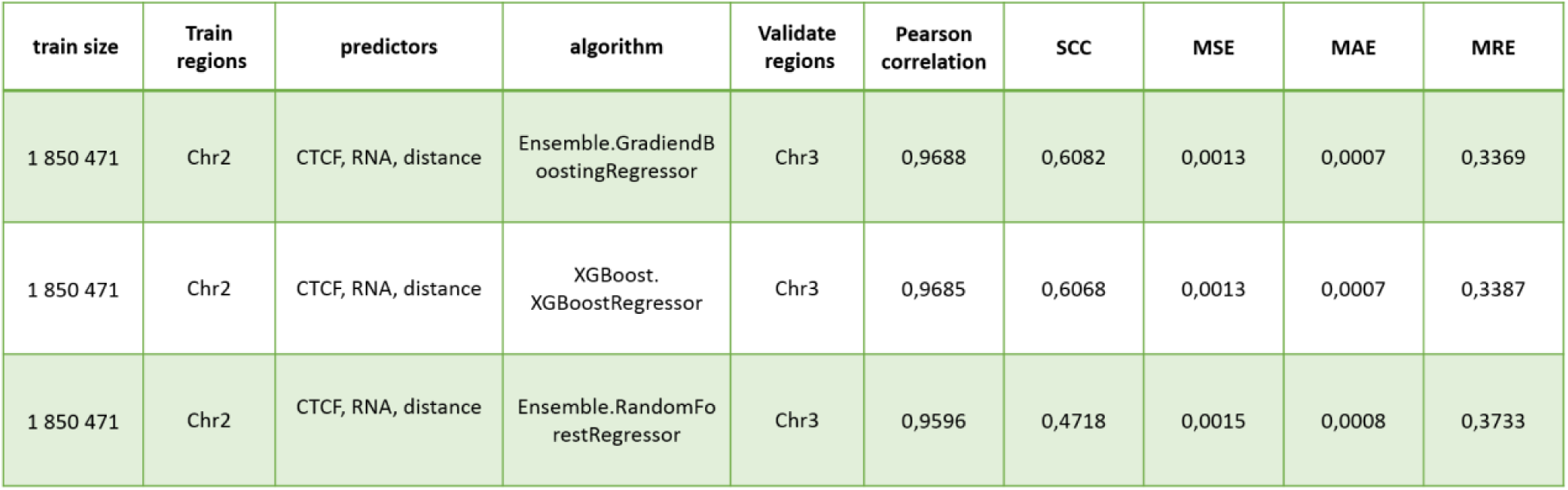
Comparison of Gradient Boosting (xgoost and sklearn implementations) and Random Forest performance.

**Supplementary Table 4.** Effect of smoothing parameter *h* on the SCC values

| train size<br>(number of<br>contacts) | Train<br>regions | predictors | Validate regions | $h$ | SCC | Pearson<br>correlation |
| --- | --- | --- | --- | --- | --- | --- |
| 25 000 | Chr10 | CTCF,<br>CTCF_orientation,<br>RNA-seq,<br>contact_distance | Chr2 | 0 | 0.572 | 0.936 |
| 25 000 | Chr10 | CTCF,<br>CTCF_orientation,<br>RNA-seq,<br>contact_distance | Chr2 | 1 | 0.638 | 0.936 |
| 25 000 | Chr10 | CTCF,<br>CTCF_orientation,<br>RNA-seq,<br>contact_distance | Chr2 | 2 | 0.651 | 0.936 |
| 25 000 | Chr10 | CTCF,<br>CTCF_orientation,<br>RNA-seq,<br>contact_distance | Chr2 | 3 | 0.657 | 0.936 |
| 25 000 | Chr10 | CTCF,<br>CTCF_orientation,<br>RNA-seq,<br>contact_distance | Chr2 | 4 | 0.659 | 0.936 |
| 25 000 | Chr10 | CTCF,<br>CTCF_orientation,<br>RNA-seq,<br>contact_distance | Chr2 | 5 | 0.660 | 0.936 |
| 25 000 | Chr10 | CTCF,<br>CTCF_orientation,<br>RNA-seq,<br>contact_distance | Chr2 | 6 | 0.659 | 0.936 |

